# Molecular MRI of Neuroinflammation in Alzheimer’s Disease using a Folate-targeted Nanoprobe

**DOI:** 10.64898/2026.10.27.761963

**Authors:** Andrew A. Badachhape, Prajwal Bhandari, Romeesa Khan, Rohan Bhavane, Renuka Menon, Parag Parekh, Prasad Admane, Esther Ngan, Mayank Srivastava, Eric A. Tanifum, Ananth V. Annapragada, Rodney M. Ritzel, Ketan B. Ghaghada

## Abstract

**Background:** Neuroinflammation, driven by activated microglia, plays a central role in the pathogenesis of Alzheimer’s disease (AD). Non-invasive imaging of neuroinflammation holds promise for advancing our understanding of disease mechanisms and evaluating anti-inflammatory therapeutic strategies. The folate receptor (FR), specifically its β isoform, has been identified as a marker of pro-inflammatory myeloid cells in peripheral inflammatory diseases; however, its association with AD neuroinflammation remains unknown. In this work, we first determined FR expression in microglia in mouse models of AD and subsequently evaluated molecular MRI (mMRI) using a folate-targeted nanoprobe for *in vivo* imaging of the folate receptor expression.

**Methods and results:** Microglial activation and FR expression in APP/PS1 and P301S transgenic mouse models of AD was measured using flow cytometry. A FR-targeted nanoprobe for mMRI was prepared by functionalizing high T1-relaxivity liposomal gadolinium nanoparticle with folic acid. Following intravenous injection, localization of the nanoprobe in the brains of aged transgenic and wild-type mice was measured using T1-weighted MRI. Post-mortem microscopy measured nanoprobe localization relative to microglia and amyloid pathology. In both mouse models, FRβ was significantly (p < 0.05) upregulated in pro-inflammatory microglia expressing TNF and IL-1β. The folate-targeted nanoprobe demonstrated MR signal enhancement in the cortical and hippocampal regions of transgenic mice but not in wild type mice; non-targeted nanoprobe control did not demonstrate signal enhancement in brain of transgenic mice. Microscopy confirmed colocalization of FR-targeted nanoprobe with IBA1-positive microglia/macrophages and amyloid plaques.

**Conclusions:** Pro-inflammatory microglia exhibit elevated levels of FR, and its measurement by non-invasive mMRI is a promising marker of neuroinflammation in AD.

## Background

Neuroinflammation is increasingly recognized as a central driver of Alzheimer’s disease (AD) pathogenesis and progression[1–3]. Microglial activation, a hallmark of neuroinflammation, contributes to early amyloid-β (Aβ) accumulation, tau pathology, and synaptic dysfunction[4–9]. Importantly, clinical studies, including those involving cognitively resilient individuals with AD pathology, have shown that microglia-induced neuroinflammation independently contributes to cognitive decline[10–12]. Non-invasive imaging of neuroinflammation could significantly enhance our ability to monitor the phenomenon in neurodegenerative diseases including AD and aid in the evaluation of therapies targeting inflammatory pathways.

The folate receptor (FR), a glycosylphosphatidylinositol-anchored cell surface protein, facilitates the intracellular delivery of folates, which are essential for one-carbon metabolism (OCM). OCM supports nucleotide synthesis (purine and thymidylate) and the methylation of homocysteine to methionine. It also serves as a key metabolic pathway that promotes the activation and function of pro-inflammatory cell states by generating critical metabolites for the epigenetic activation of IL-1β–mediated inflammatory responses[13]. The β isoform of folate receptor (FRβ) has emerged as a marker of pro-inflammatory myeloid cells in multiple inflammatory diseases, including encephalomyelitis and multiple sclerosis[14–19]; however, its association with AD neuroinflammation remains unknown.

In this study, we first performed functional phenotyping of microglia using flow cytometry to assess the expression of folate receptor isoforms (FRα and FRβ) in two transgenic Alzheimer’s disease (AD) mouse models. Subsequently, we developed a novel high T1-relaxive folate-targeted nanoprobe and evaluated its performance for molecular MRI (mMRI) of neuroinflammation. Our findings demonstrate that FR is upregulated in pro-inflammatory microglia and that the folate-targeted nanoprobe enables mMRI-based detection of neuroinflammation in AD mouse models.

## Materials and Methods

### Animal models

All animal studies were performed under a protocol approved by the Institutional Animal Care and Use Committee (IACUC). Studies were performed in two transgenic (TG) mouse models of Alzheimer’s disease: APPswe/PSEN1dE9 (APP/PS1) (Strain #034832-JAX, The Jackson laboratory, ME, USA) double transgenic mouse model of amyloid pathology and early-onset Alzheimer’s disease, and B6;C3-Tg(Prnp-MAPT*P301S)PS19Vle/J (P301S) (Strain # 008169, The Jackson laboratory, ME, USA) transgenic mouse model of tau pathology which express the P301S mutant form of human microtubule-associated protein tau (MAPT). APP/PS1 transgenic mice develop amyloid plaques in the brain starting at 6–7 months of age[20,21]. Tau P301S transgenic mice develop neurofibrillary tangles by 5 months of age[22,23]. Transgenic mice of both strains have previously been shown to exhibit microglial activation and neuroinflammation[20,22,24]. In this work, we used 11–18-month-old APP/PS1 mice and 9 – 11-month-old Tau P301S mice, ages at which frank pathology is observed in each strain. All studies of transgenic mice included age-matched wild type siblings as controls.

### *In vivo* MRI

*In vivo* imaging was performed on a 1T permanent magnet scanner (M7 system, Aspect Imaging, Shoham, Israel) using methods described previously[25,26]. Mice were sedated with 2-3% isoflurane and placed on a custom, MRI-safe bed with an integrated head-cone for continuous delivery of anesthesia. Animals underwent pre-contrast MRI followed by intravenous administration of either folate-targeted nanoprobe (0.2 mmol Gd/kg) or non-targeted control nanoprobe (0.2 mmol Gd/kg). Post-contrast scans were performed four days after administration of nanoprobe. A T1-weighted spin echo (T1w-SE) sequence with the following parameters was used for image acquisition: TR = 600 ms, TE = 11.5 ms, matrix = 192×192, FOV = 30 mm, slices = 16, slice thickness = 1.2 mm, NEX = 2. Each animal underwent two T1w-SE scans, and the data were externally averaged to reduce motion artifact. Data processing was performed in MATLAB (version R2022b, Mathwords, Natick, MA) and MR signal intensity analysis was performed in OsiriX (version 5.8.5, 64-bit, Pixmeo SARL, Geneva, Switzerland).

MR signal intensity (SI) was calculated by segmentation of a 3D region-of-interest (ROI) in the cortical regions of three contiguous central slices in each image. SI was compared in pre-contrast, baseline scans and 4-day post-contrast scans. MR signal enhancement was calculated as (SI_post_ – SI_pre_)/SI_pre_ and expressed as a percent increase.

### Preparation of nanoprobes

The folate-targeted nanoprobe (FNP) is based on a high T1 relaxive liposomal-Gd nanoparticle platform that has previously been used as the basis of targeted contrast agents for mMRI of amyloid and tau pathology in Alzheimer’s disease [25,26]. Preparation followed methods analogous to those described previously [25,26]. Briefly, L-α-phosphatidylcholine, hydrogenated (Hydro Soy PC; HSPC), cholesterol, 1,2-Distearoyl-sn-glycero-3-phosphorylethanolamine gadoterate meglumine (Gd(III)-DSPE-DOTA), 1,2-distearoyl-sn-glycero-3-phosphoethanolamine-N-[methoxy(polyethylene glycol)-2000] (DSPE-mPEG2000), 1,2-distearoyl-sn-glycero-3-phosphoethanolamine-N-[folate(polyethylene glycol)-3400] (DSPE-PEG3400-folate) and lis-rhodamine-DHPE (fluorescent tag lipid) were dissolved in ethanol at molar ratio of 31.4:40:25:3:0.5:0.1. A non-targeted nanoprobe (NT) without folate conjugate was prepared and used as control formulation for *in vivo* studies. The ethanolic solution of lipids was hydrated with saline solution (150 mM) at 65 °C for 30 minutes, and then sequentially extruded in a 10ml Lipex extruder using a 400 nm Whatman Nuclepore polycarbonate filter (4 passes) followed by a 200 nm (4 passes) and finally through 100 nm filters (8 passes). The resultant suspension was diafiltered against 150 mM saline solution and concentrated using a *750 kDa* cutoff cross-flow diafiltration cartridge. Gd and P concentrations were measured using inductively coupled plasma atomic emission spectroscopy (ICP-AES). Particle size distribution, hydrodynamic diameter and polydispersity index of nanoprobes were measured by dynamic light scattering.

### Flow cytometry

Mice were perfused i.v. with 40 mL of cold phosphate-buffered saline. The whole brain was excised and then halved along the interhemispheric fissure [27]. The left hemisphere was isolated after removing the olfactory bulb and cerebellum. Using a 5 mL Luer-Lok syringe (Becton Dickinson) plunger, the brain tissue was mechanically digested through a 70-μm cell strainer (CellTreat) cap placed on a 50 mL conical tube (Falcon) containing complete Roswell Park Memorial Institute (RPMI) 1640 (Invitrogen, Cat# 22400105) medium, and immediately placed on ice. These samples were enzymatically digested in collagenase/dispase (1 mg/ml, Roche Diagnostics, Cat# 10269638,001), papain (5U/ml, Worthington Biochemical, Cat# <u>LS003119</u>), 0.5 M EDTA (1:1000, Invitrogen, Cat# 15575020), and DNAse I (10 mg/ml, Roche Diagnostics, Cat# 10104159001) for 1 h at 37 °C on a shaking incubator (200 rpm). The cell suspension was washed twice with RPMI, filtered through a 70-μm filter, and RPMI was added to a final volume of 5 mL/hemisphere and kept on ice. Cells were then transferred into FACS tubes and washed with FACS buffer. Brain cell suspensions were incubated with Fc block (Anti-CD16/32, BioLegend, Cat#101302) for 10 minutes on ice prior to staining for the following surface antigens: CD45-eF450 (30-F11, eBioscience, Cat# 48–0451−82), CD11b-APCeF750 (M1/70, BioLegend, Cat#101262), Ly6C-AF700 (HK1.4, BioLegend, Cat#128024). The fixable viability dye Zombie UV was used for live/dead discrimination (BioLegend, Cat#423108). Cells were then washed in FACS buffer, fixed in 2% paraformaldehyde for 10 min, and washed once more prior to adding 500 μl FACS buffer. Due to rapid internalization, cell surface and intracellular staining for folate receptor α (FRα)-FITC (polyclonal, Fabgennix, Cat#FOLR1-FITC) and folate receptor β (FRβ)-APC and -PE (10/FR2, BioLegend, Cat#153306 and 153304, respectively) was performed to measure total FR expression. Intracellular staining for Lamp1-PerCPCy5.5 (1D4B, BioLegend, Cat#121626) was performed using Cytofix/Cytoperm Fixation/Permeabilization Kit (BD Biosciences, Cat#554714) according to manufacturer’s instructions and as described previously [28]. Cytokine staining for TNF-PECy7 (BioLegend, Cat#506324) and IL-1β-PerCPeF710 (eBioscience, Cat#46–7114-82) was performed after 2 h incubation with Brefeldin A (BioLegend, Cat#420601) at 37°C followed by fixation/permeabilization. Cells were loaded for 20 min in a 37°C water bath, washed three times with FACS buffer (without NaAz), and then stained for surface markers including viability stain.

Data were acquired on a Beckman-Coulter CytoFlex LX cytometer using CytExpert v2.6 software and then analyzed using FlowJo (Treestar Inc.). At least 5–10 million events were collected for each sample. CountBright Absolute Counting Beads (20 μL/sample, ThermoFisher, Cat#C36950) were used to estimate cell counts per the manufacturer’s instructions. Data were expressed as counts/hemisphere. Leukocytes were first gated using a splenocyte reference (SSC-A vs FSC-A). Singlets were gated (FSC-H vs FSC-W), and live cells were gated based on Zombie Aqua exclusion (SSC-A vs Zombie UV). Resident microglia were identified as CD45^int^CD11b^+^Ly6C^−^, whereas peripheral leukocytes were distinguished as CD45^hi^CD11b^+^ myeloid cells or CD45^hi^CD11b^−^ lymphocytes. Cell type–matched fluorescence minus one (FMO) controls were used to determine the positivity of each antibody.

### Tissue preparation and microscopy

Following euthanasia, transgenic (*APP/PSEN1*) and wild-type mice underwent transcardial perfusion to ensure thorough tissue fixation and vascular clearance. The perfusion protocol commenced with a 0.9% saline solution to remove residual blood, followed by a 4% paraformaldehyde (PFA) solution in 0.1 M phosphate-buffered saline (PBS, pH 7.4) to preserve tissue morphology and antigenicity. Post-perfusion, brains were carefully extracted and immersed in 4% PFA at 4°C for 48 hours. Subsequently, tissues were transferred to a 30% sucrose solution in PBS. The sucrose solution was changed daily until the brains sank, indicating complete cryoprotection.

Following cryoprotection, brains were embedded in Optimal Cutting Temperature (OCT) compound and flash-frozen using dry ice. Once solidified, the tissue blocks were stored at −80°C until sectioning. For sectioning, brains were equilibrated to approximately −20°C and cut into 50 µm coronal sections using a cryostat. To unmask antigens, tissue sections underwent microwave-assisted antigen retrieval. Sections were incubated in a preheated citrate buffer (10 mM citric acid, 0.05% Tween 20, pH 6.0) at an intermediate power setting for two cycles of 5 minutes each, with a 1-minute interval between cycles. After retrieval, sections were allowed to cool at room temperature for 15 minutes.

Immunofluorescence staining was performed by incubating sections overnight at 4°C with primary antibodies: a 1:200 dilution of β-amyloid antibody (Alexa Fluor® 647 anti-β-Amyloid, 17-24 Antibody, Biolegend), and a 1:200 dilution of IBA1 antibody (Iba1/AIF-1 (E4O4W), Rabbit mAb #17198) for microglia. Following primary antibody incubation, sections were washed with PBS containing 0.1% Tween 20 (PBST) and incubated for 2 hours at room temperature with appropriate secondary antibodies at a 1:1000 dilution. Nuclear counterstaining was achieved using DAPI after additional PBST washes. Finally, sections were mounted using Fluoromount G (Electron Microscopy Sciences, 17984) and visualized using an Olympus Fluoview LV100 confocal microscope at 40x magnification. Single-channel IBA-1 and FNP signals were quantified using Imaris software Version 10.2, and co-localization of fluorescence signals was analyzed with Cytseer software (Version 2.8.1.0) using the Pearson correlation coefficient (PCC) to assess spatial overlap.

### Data and statistical analysis

Statistical analyses were performed using GraphPad Prism 10.0. Comparison of MR signal change between pre-contrast and post-contrast scans was performed using a Wilcoxon rank-sum test to test for pairwise group differences between three cohorts: 1) Tg mice tested with FNP(n=6 APP/PS1, n=9 P301S), 2) WT animals tested with FNP (n=6 APP/PS1 WT littermates, n=6 P301S WT littermates), and 3) Tg mice tested with control, non-targeted nanoprobes (n=5 APP/PS1, n=4 P301S). Group comparison of flow cytometric data and fluorescence microscopy data was performed using Student’s T-test. Values of p ≤ 0.05 were nominally considered significant for all statistical analysis.

## Results

Flow cytometry revealed differential microglial activation and folate receptor (FR) expression in the brains of APP/PS1 and P301S mouse models of AD. In the APP/PS1 model, Tg animals exhibited significantly elevated levels of CD45^int^CD11b^+^Ly6C^−^ microglia compared to WT controls (**Fig 1A**), indicating ongoing microgliosis. Microglial production of the pro-inflammatory cytokine, tumor necrosis factor (TNF), was significantly higher in Tg mice compared to WT mice (**Fig 1B**), suggesting heightened microglial activation.

**Figure 1.**
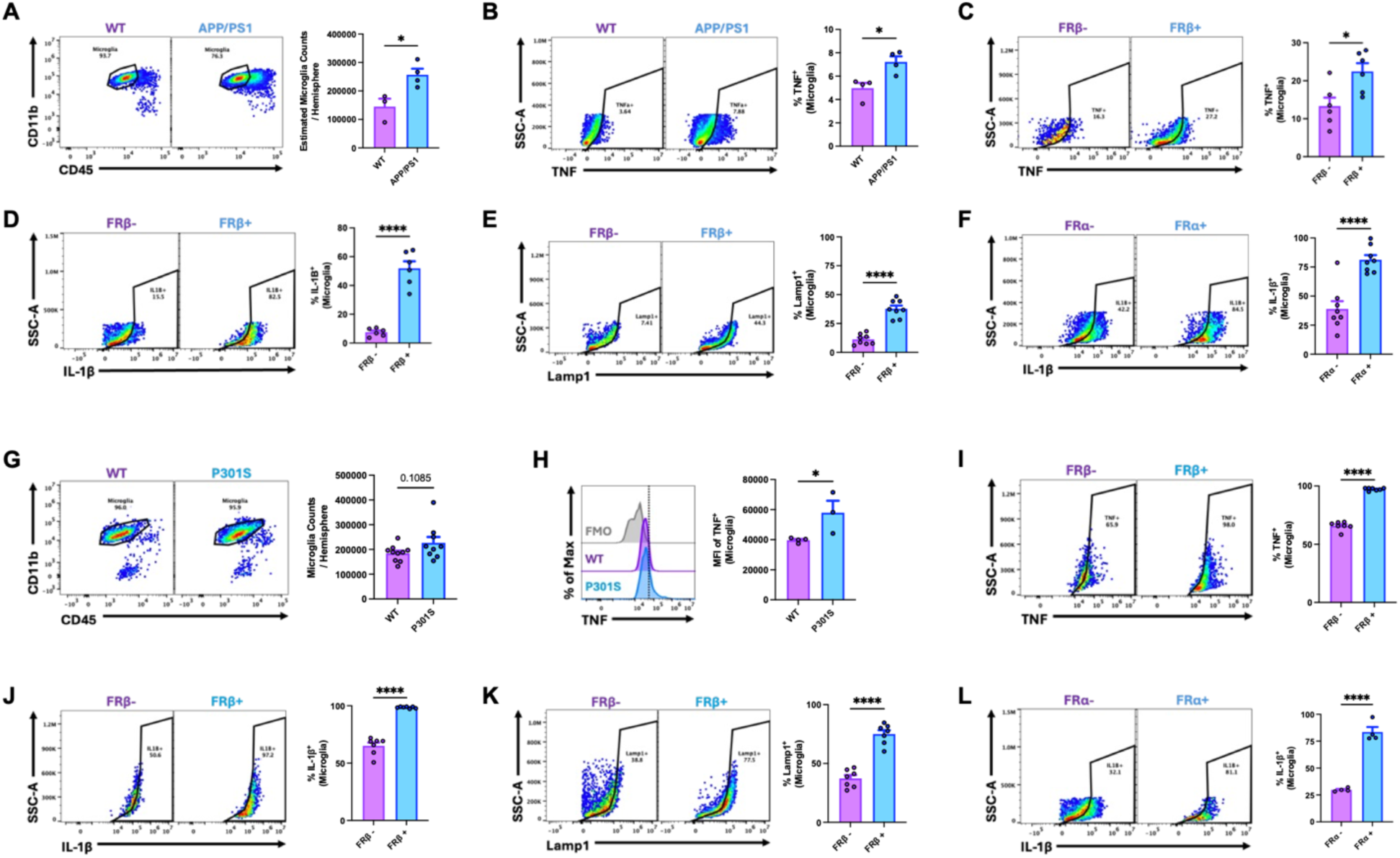
High expression of folate receptor constitutes a marker of pro-inflammatory microglia. Flow cytometric evaluation of microglial activation status and folate receptor expression in APP/PS1 and P301S mice. <u>APP/PS1:</u> (A) Transgenic (Tg) mice demonstrated higher levels of CD45^int^CD11b^+^Ly6C^−^ microglia compared to wildtype (WT) control mice. (B) Microglial production of pro-inflammatory cytokine, TNF, was significantly higher in Tg mice compared to WT mice. (C) The percentage of TNF^+^ microglia was significantly higher in FRβ^+^ microglia compared to FRβ^−^ microglia. (D) The percentage of IL-1β^+^ microglia was significantly higher in FRβ^+^ microglia compared to FRβ^−^ microglia. (E) The percentage of Lamp1-positive microglia was significantly higher in FRβ^+^ microglia compared to FRβ^−^ microglia. (F) The percentage of IL-1β^+^ microglia was significantly higher in FR*a*^+^ microglia compared to FR*a*^−^ microglia. <u>P301S:</u> (G) Microglial counts were not significantly different in Tg and WT mice. (H) Microglial production of pro-inflammatory cytokine, TNF, was significantly higher in Tg mice compared to WT mice. (I) The percentage of TNF^+^ microglia was significantly higher in FRβ^+^ microglia compared to FRβ^−^ microglia. (J) FRβ^+^ microglia produced higher level of pro-inflammatory cytokine, IL-1β, compared to FRβ^−^ microglia. (K) The percentage of Lamp1-positive microglia was significantly higher in FRβ^+^ microglia compared to FRβ^−^ microglia. (L) FR*a*^+^ microglia produced higher level of pro-inflammatory cytokine, IL-1β, compared to FR*a*^−^ microglia.

To characterize the inflammatory phenotype in FR-expressing microglia, cytokine production in FR-positive versus FR-negative microglia was compared. FRβ^+^ microglia exhibited significantly higher expression of TNF and IL-1β than FRβ^−^ microglia (**Fig 1C, D**), and also showed elevated levels of Lamp-1 (**Fig 1E**), indicating microglial activation associated with higher phagocytic/lysosomal activity. Similarly, FR*a*^+^ microglia demonstrated significantly higher IL-1β expression compared to FR*a*^−^ microglia. (**Fig 1F**).

In the P301S model, levels of CD45^int^CD11b^+^Ly6C^−^ microglia were not significantly different between Tg animals and WT controls (**Fig 1G**). Microglial production of TNF was significantly higher in Tg mice compared to WT mice (**Fig 1B**), suggesting a heightened activation state of microglia (**Fig 1H**). FRβ^+^ microglia in these mice expressed significantly higher levels of TNF, IL-1β and Lamp-1 than FRβ^−^ microglia (**Fig 1I, J, K**), and FR*a*^+^ microglia similarly exhibited elevated IL-1β expression (**Fig 1L**). These findings collectively suggest that FRα and FRβ are markers of a pro-inflammatory and dysfunctional microglial phenotype in both APP/PS1 and P301S models.

Physicochemical characterization of FNP (**Fig 2**) indicated a mean particle size of ∼100 nm, a polydispersity index of 0.15 and an estimated 36,600 gadolinium atoms and 760 folate ligands per nanoparticle. Folate receptor and neuroinflammation levels in the APP/PS1 and P301S models were both studied in vivo by mMRI using this nanoprobe. Pre-contrast T1-weighted spin echo (T1w-SE) scans were acquired, followed by intravenous administration of the FNP. Non-targeted nanoprobe (NT) was used as a control agent for testing in transgenic mice. Delayed imaging was conducted four days post-injection to allow clearance of circulating probe thus minimizing blood pool signal, based on our previous studies[25,26] (**Fig. 3A**). In both mouse models of AD, post-contrast images demonstrated MR signal enhancement in cortical and hippocampal regions of Tg mice administered FNP, while WT mice administered FNP showed no enhancement (**Fig. 3B, C**). Transgenic mice administered NT showed no signal enhancement. Quantitative analysis confirmed significantly higher MR signal in Tg brains compared to WT controls in mice administered FNP and Tg mice administered NT (**Fig. 3B, C**). No significant differences were seen between Tg and WT mice administered NT and FNP, respectively.

**Figure 2.**
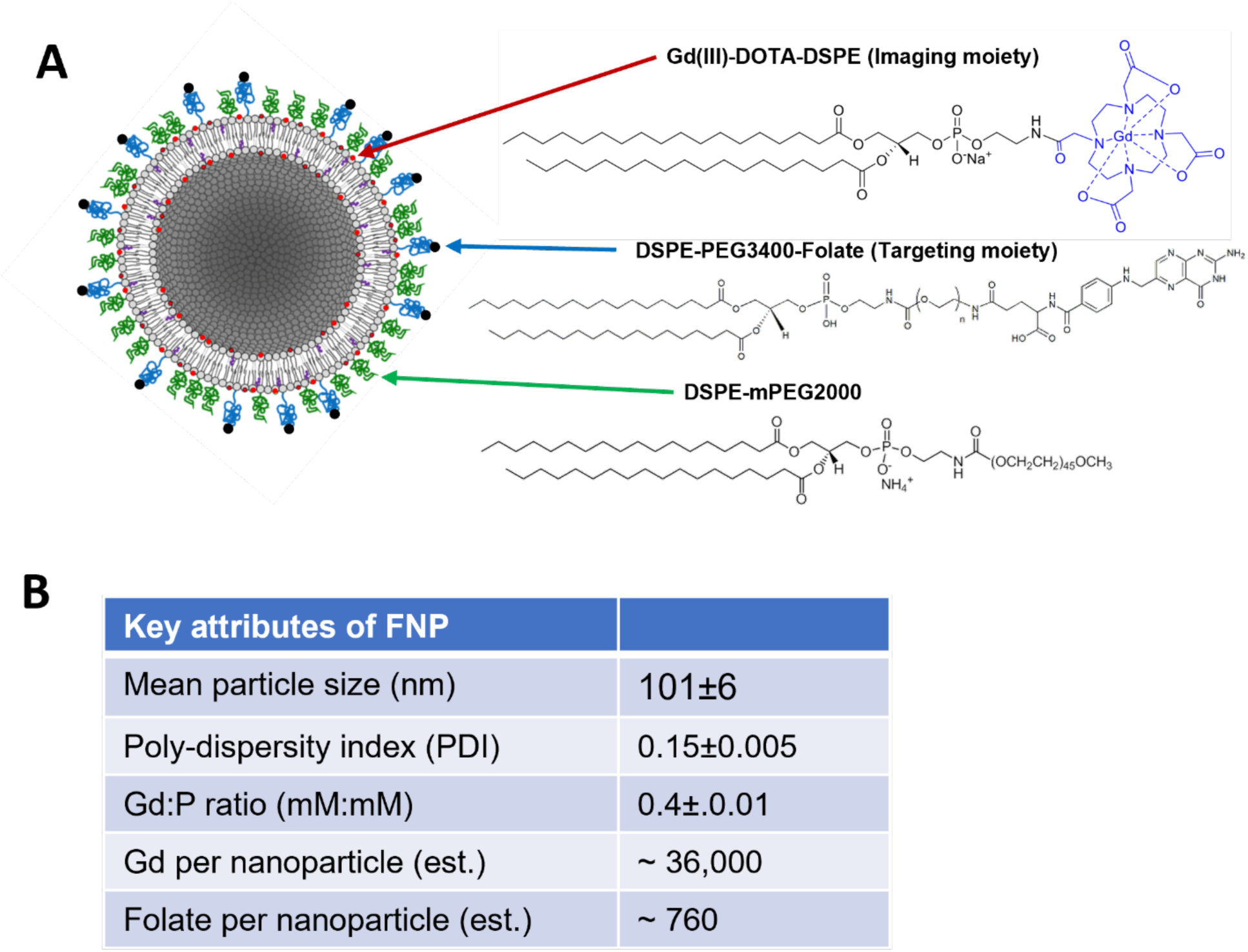
Folate-targeted nanoprobe (FNP). **(A)** Schematic of folate-targeted liposomal-Gd nanoprobe showing structures of key moieties. **(B)** Key attributes of FNP.

**Figure 3.**
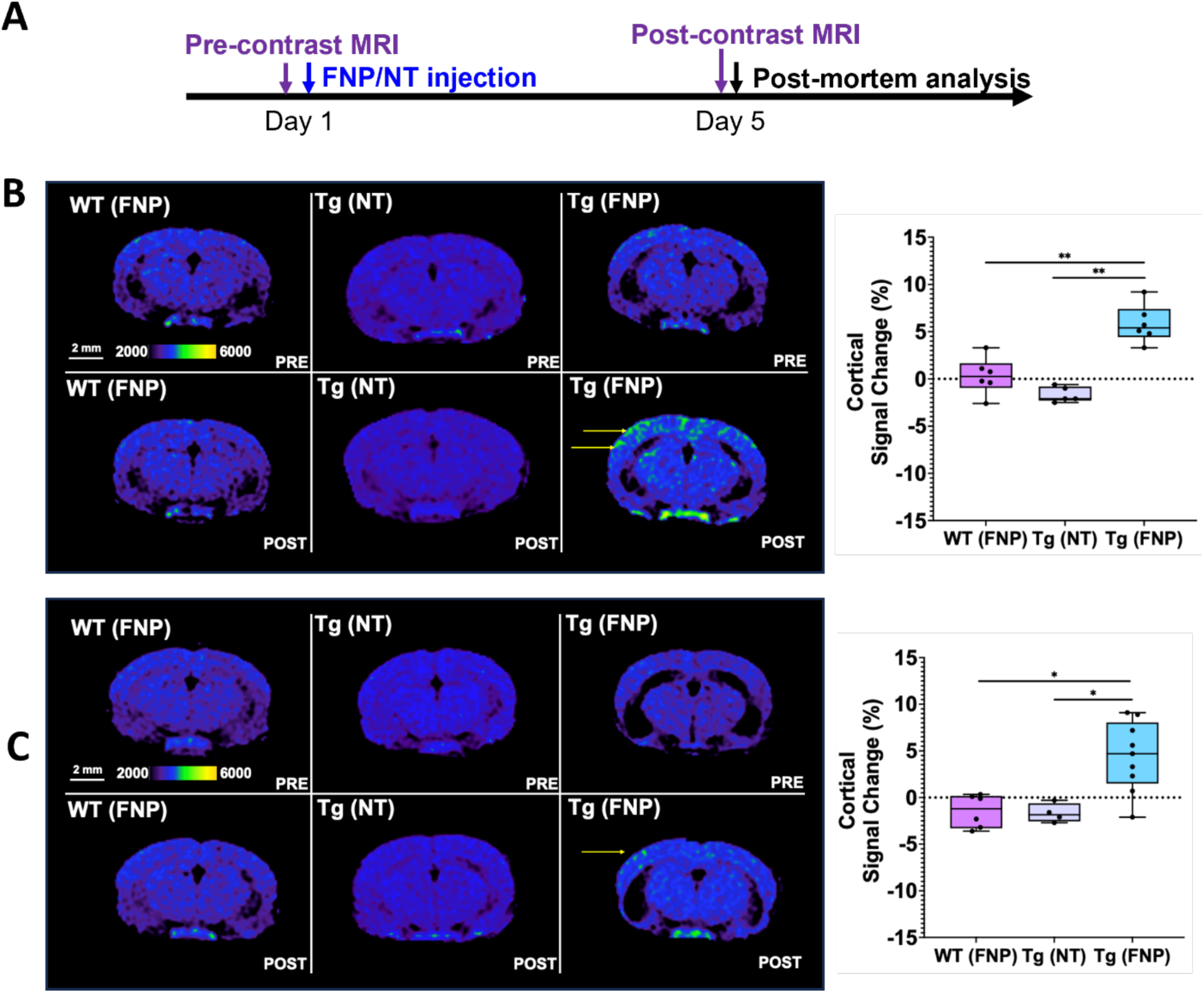
FNP enables *in vivo* MRI of neuroinflammation in Alzheimer’s disease (AD) mice. **(A)** Experimental timeline for testing FNP in mouse models of AD. Pseudo-colored T1-weighted spin-echo (T1w-SE) MR images in axial plane and quantitative analysis of MR signal changes in **(B)** APP/PS1 and **(C)** Tau P301S mouse models. Cortical regions demonstrate significant signal enhancement in both sets of Tg mice tested with FNP relative to both WT mice tested with FNP and Tg mice tested with NT (*p<0.05; **p<0.005). Scale bar represents 2 mm. Legend: FNP, folate-targeted nanoprobe; NT, non-targeted nanoprobe; Tg, transgenic; WT, wild type.

Histological characterization of post-mortem brain tissue from APP/PS1 mice revealed strong colocalization of FNP with IBA1-positive microglia/macrophages in regions of amyloid plaque deposition in APP/PS1 Tg mice (**Fig. 4A**). Fluorescence intensity of FNP was significantly higher in Tg brains than in WT (**Fig. 4B**), corroborating the MRI findings. Co-localization analysis showed a strong Pearson’s correlation coefficient (PCC) between FNP signal and both microglia/macrophages and amyloid pathology in Tg mice (**Fig. 4C, D**).

**Figure 4.**
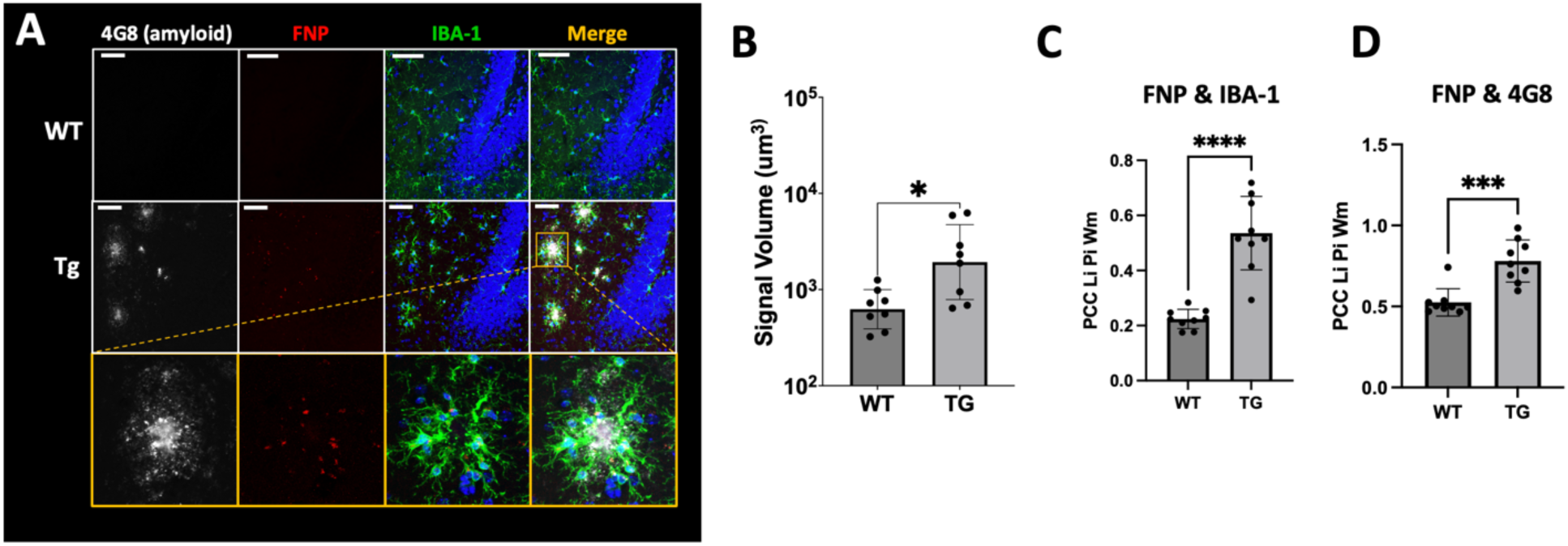
FNP co-localizes with microglia-like cells in regions of AD pathology. **(A)** Immunofluorescence analysis of brain sections from APP/PS1 Tg and WT mice show accumulation of fluorescently labeled FNP in regions of amyloid plaque (4G8 staining) and co-localization with microglia/macrophage (large soma in IBA-1 staining). Scale bar represents 200 μm. **(B)** Quantitative analysis of fluorescence images demonstrates significantly higher (p<0.05) folate-targeted nanoprobe (FNP) signal in Tg brains compared to WT brains. Pearson correlation coefficient (PCC) analysis shows stronger co-localization of **(C)** FNP with IBA-1 positive microglia/macrophage, and **(D)** FNP with amyloid plaques, in brains of Tg mice compared to WT mice. Legend: FNP, folate-targeted nanoprobe; Tg, transgenic; WT, wild type.

## Discussion

In this study, we demonstrate that neuroinflammation in mouse models of Alzheimer’s disease (AD) is associated with folate receptor (FR) over-expression and microglial activation. FR-expressing microglia exhibited a pro-inflammatory phenotype in both amyloid and tau models of AD. Furthermore, a FR-targeted nanoprobe for molecular MRI (mMRI) localized to IBA1-positive microglia/macrophages in regions of amyloid plaque deposition, resulting in MR signal enhancement in the cortical and hippocampal regions of transgenic mice but not in wild type mice. These findings link FR-expressing, pro-inflammatory microglia to the sites of AD pathology.

Prior studies have shown that high expression of the FRβ isoform is a marker of pro-inflammatory myeloid cell phenotype in multiple inflammatory diseases[14,15], however, its role in AD remains unexplored. Using flow cytometry-based co-expression analysis, we demonstrate that pro-inflammatory microglia in AD mouse models, that exhibit elevated TNF and IL-1β expression, overexpress both FRα and FRβ isoforms. These findings position FR as a promising marker of activated microglia in AD.

Leveraging a highly T1-relaxive liposomal-Gd nanoprobe platform, previously validated for amyloid and tau mMRI, we developed a folate-targeted nanoprobe and tested in APP/PS1 and P301S mouse models of AD. Aged transgenic mice, known to exhibit neuroinflammation, were used for in vivo mMRI studies. Our data show that the folate-targeted nanoprobe penetrates the brain, localizes to microglia, and enables high-resolution *in vivo* imaging of neuroinflammation. These results establish proof of concept for FR-targeted mMRI for detecting and monitoring neuroinflammation in AD.

This work has several limitations. A deeper microglial phenotyping is necessary to assess the co-expression of FR isoforms alongside other pro- and anti-inflammatory markers, as well as additional functional and metabolic indicators. Such comprehensive profiling would enhance our understanding of microglial FR expression during neuroinflammation in AD, a condition known to involve altered cellular metabolism. The relationship between neuroinflammatory burden and probe sensitivity also warrants further investigation. Interpretation of the FRα findings requires further investigation. Unlike FRβ, which is well established in activated monocyte/macrophage lineage cells, FRα expression has traditionally been associated with epithelial cell populations, particularly the choroid plexus epithelium[29]. Furthermore, the polyclonal anti-FRα antibody used in this study may be susceptible to cross-reactivity. Therefore, additional studies using orthogonal validation approaches will be required to confirm FRα expression and define its biological significance in AD-associated microglia. Moreover, the potential uptake of the targeted nanoprobe by other brain cell types should be explored. Mechanistically, the drivers of FR upregulation during microglial activation remain undefined. Folate deficiency has been shown to increase homocysteine levels, a known risk factor for AD and dementia[30,31]. Preclinical studies suggest that folate deficiency mediates neuroinflammation and cognitive impairment via activation of the NF-ĸB signaling pathway[32]. Furthermore, folate deficiency has been associated with microglial activation, while folate supplementation has been shown to restore microglial homeostasis and exert neuroprotective effects[33]. FRs are upregulated in folate deficiency and downregulated in folate repletion, reflecting the dynamic regulation of folate transport in response to cellular demand. Upregulation of FR can also be triggered by cell activation due to increased folate demand in one-carbon metabolism (OCM), which is required for the production of metabolites that drive IL-1β–mediated inflammatory responses[13,34]. Evidence from peripheral models supports roles for homocysteine-mediated translational regulation and metabolic reprogramming[35], processes likely seen in AD microglia, which shift toward glycolysis and rely on OCM to sustain pro-inflammatory states. These pathways may underlie the observed FR upregulation and support its potential as both a biomarker and a therapeutic target.

## Conclusion

This study identifies over-expression of folate receptor as a potential marker of pro-inflammatory microglia in Alzheimer’s disease. Furthermore, it demonstrates the feasibility of molecular MRI for imaging neuroinflammation in Alzheimer’s disease using a folate receptor-targeted nanoprobe.

## Declarations

## Ethics approval and consent to participate

All animal studies were approved by the Baylor College of Medicine Animal Care and Use Committee.

## Consent for publication

All authors have consented for the publication of manuscript.

## Availability of data and materials

Data supporting the findings reported in this article are available from the corresponding author on reasonable request.

## Authors’ contributions

Andrew A. Badachhape: Conceptualization; investigation; writing-original draft; writing-review and editing. Prajwal B handari: Investigation; writing-original draft; writing-review and editing. Romeesa Khan: Investigation; writing-review and editing. Rohan Bhavane: Investigation; writing-review and editing. Renuka Menon: Investigation; writing-review and editing. Parag Parekh: Investigation; writing-review and editing. Prasad Admane: Investigation; writing-review and editing. Esther Ngan: Investigation; writing-review and editing. Mayank Srivastava: Investigation; writing-review and editing. Eric A Tanifum: Investigation; writing-review and editing. Ananth V Annapragada: Writing-review and editing, supervision. Rodney Ritzel: Conceptualization; investigation; writing-original draft; supervision, writing-review and editing. Ketan Ghaghada: Conceptualization; investigation; writing-original draft; supervision, writing-review and editing. All authors have read and approved the final version of the manuscript.

